# Potential microenvironment of SARS-CoV-2 infection in airway epithelial cells revealed by Human Protein Atlas database analysis

**DOI:** 10.1101/2020.04.16.045799

**Authors:** Ling Leng, Jie Ma, Leike Zhang, Wei Li, Lei Zhao, Yunping Zhu, Zhihong Wu, Ruiyuan Cao, Wu Zhong

## Abstract

The outbreak of COVID-19 has caused serious epidemic events in China and other countries. With the rapid spread of COVID-19, it is urgent to explore the pathogenesis of this novel coronavirus. However, the foundational research of COVID-19 is very weak. Although angiotensin converting enzyme 2 (ACE2) is the reported receptor of SARS-CoV-2, information about SARS-CoV-2 invading airway epithelial cells is very limited. Based on the analysis of the Human Protein Atlas database, we compared the virus-related receptors of epithelial-derived cells from different organs and found potential key molecules in the local microenvironment for SARS-CoV-2 entering airway epithelial cells. In addition, we found that these proteins were associated with virus reactive proteins in host airway epithelial cells, which may promote the activation of the immune system and the release of inflammatory factors. Our findings provide a new research direction for understanding the potential microenvironment required by SARS-CoV-2 infection in airway epithelial, which may assist in the discovery of potential drug targets against SARS-CoV-2 infection.

## Introduction

The outbreak of COVID-19 pneumonia was declared a “public health emergency of international concern” by the World Health Organization (WHO). On January 7, 2020, Chinese scientists isolated a new type of coronavirus named SARS-CoV-2 from an infected patient. SARS-CoV-2 causes serious respiratory diseases and is transmitted by human-to-human contact [1]. However, the pathogenesis of this novel pathogen and any potential drug targets remain to be elucidated. The clinical symptoms of SARS-CoV-2 resembles those of SARS-CoV [2], and the genome of SARS-CoV-2 represents good homology with SARS-CoV [3], with 79.5% sequence consistency the two coronaviruses [4]. SARS-CoV mainly replicates in the lower respiratory tract, resulting in diffuse alveolar damage [5]. The tissue tropism of SARS-CoV-2 also is the lower respiratory tract [6, 7]. In addition, it was reported that cytopathic effects were observed 96 h after inoculation of SARS-CoV-2 on the surface layers of human airway epithelial cells (EpCs), and a lack of cilium beating was observed after virus infection [8]. However, it was found that Huh7 epithelial-like cells from the liver that are similar to airway EpCs produced no specific cytopathic effects after six days of virus infection [8]. In addition, a recent study has shown that an epithelial cell line from cervical cancer (HeLa) was also insensitive to SARS-CoV-2 infection [4]. These studies suggest that cells from different epithelial sources have different susceptibility to the SARS-CoV-2.

Coronaviruses are enveloped and require a series of interactions with their host cells during virus entry, depending on tissue and cell-type specificities (e.g., receptor and microenvironment) in addition to strain and species [9]. Although some microorganisms may display a commensal relationship with EpCs [10], different microorganisms will specifically attack the EpCs of different organs. For example, SARS-CoV mainly binds to cells expressing angiotensin-converting enzyme 2 (ACE2), which is one of the major receptors of SARS-CoV when entering cells [11]. To date, more studies have confirmed that ACE2 is also the receptor of SARS-CoV-2 infection [4]; however, ACE2 displays high expression in the kidney, bile duct, and testis with low expression in the airways and lungs [12–14]. Lindskog, et al also emphasized that the expression of ACE2 in human respiratory system is limited, and questioned the role of ACE2 for infection in human lungs [15]. Therefore, we wonder whether there are some key local microenvironment proteins specifically expressed on the surface of airway EpC that makes the virus prefer airway EpCs. In some cases, additional cell surface molecules or co-receptors are required for sufficient viral entry into host cells. It remains unclear whether these local environments with specific components are related to the susceptibility of different cell types to SARS-CoV-2.

To study the specificity of airway EpCs for SARS-CoV-2 infection, we compared host cell receptors and virus infection-related host proteins expressed in epithelial or epithelial-derived cells from different organs and found a series of specific proteins and key receptors of airway EpCs related to virus infection. These proteins may be beneficial to the design of potential drug targets against SARS-CoV-2 infection.

## Materials and methods

### Data sources

Generally, the expression profiles of human genes on the protein level were derived from the Human Protein Atlas (HPA, https://www.proteinatlas.org/) [16]. The virus-related biological and functional gene ontology (GO) annotation terms were derived from the Gene Ontology knowledgebase (http://geneontology.org/, released on 2020-01-01, containing 44,700 GO terms) [17]. All human protein annotations were downloaded from the Swiss-Prot database of UniProt (https://www.uniprot.org/) [18].

### Data analysis

The overall analysis pipeline is shown in Figure 1A. First, all GO terms that related to the virus-host functions were collected from the GO-obo file. Then, after manually checking, only those terms related to host-virus receptor and key biological processes were taken (Table S2), and the human proteins involved in these particular GO terms were retrieved from UniProt database. After that, the protein expression levels in epithelial and epithelial derived cells from different organs were derived from the HPA and mapped to these proteins.

**Figure 1.**
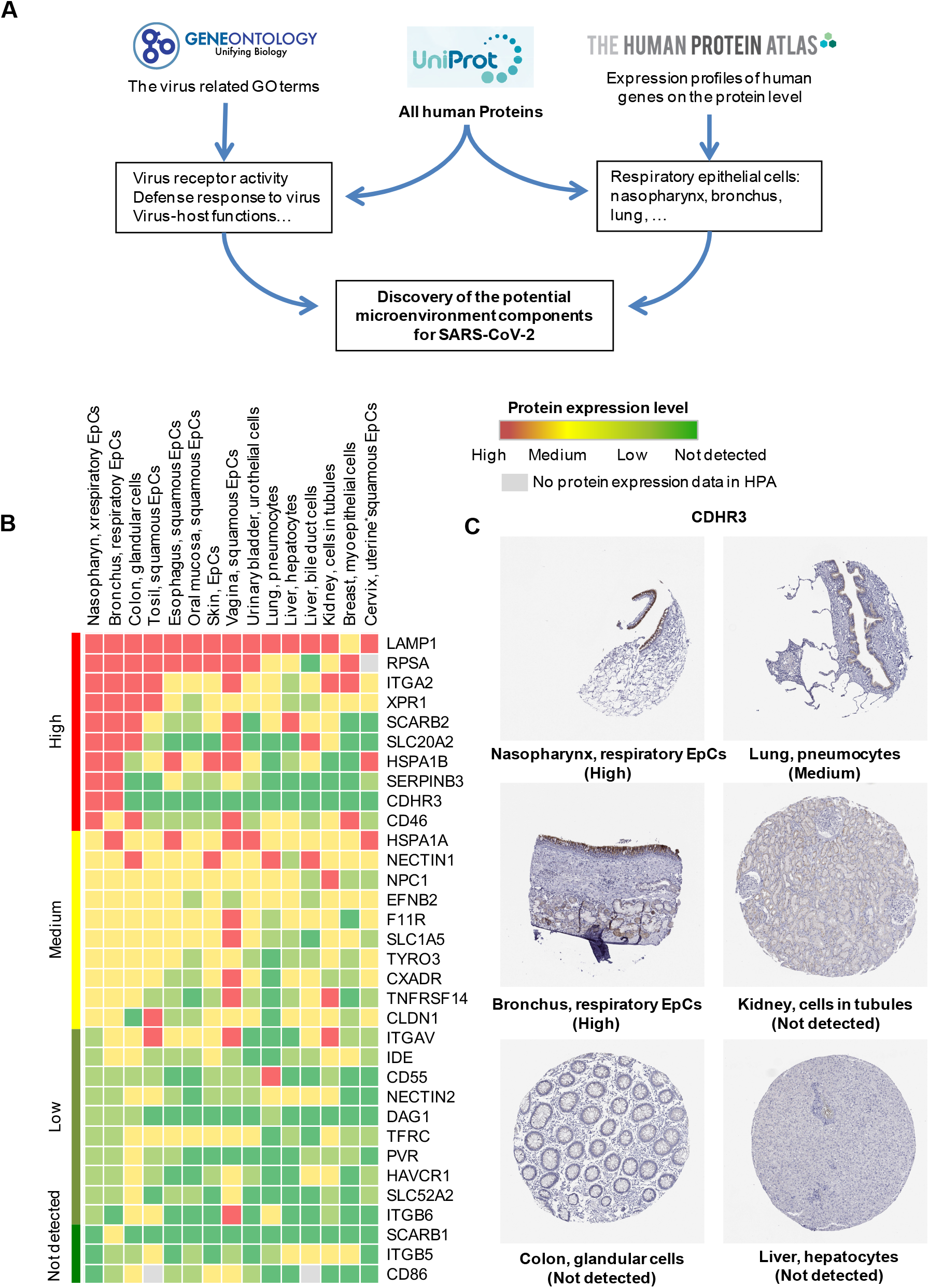
Expressions of virus microenvironment components in epithelial-derived cells from different organs. (A) Schematic illustration of data analysis strategies. (B) Heat map analysis of virus microenvironment components in epithelial-derived cells from different organs. The boxes with red, yellow, green, and dark green represent the expression levels of receptors with high, medium, low, and not detected values, respectively. (C) Immunohistochemistry analysis of CDHR3 in epithelial or epithelial derived cells of different organs. Expression profiles of all receptors are shown in Table S1, and the staining data and pictures are downloaded from “The Tissue Atlas” of HPA (https://www.proteinatlas.org/ENSG00000128536-CDHR3/tissue).

### Data visualization

Heap map views of protein expression levels were displayed using conditional formatting of Microsoft Excel spreadsheets. All protein-protein interactions were retrieved from the STRING database [19], and the protein-protein interaction network was built using Cytoscape software (version 3.7.1) [20].

## Results and discussion

### Expression of virus-associated proteins on epithelial cells is organ-specific

First, ACE2 was found highly expressed in intestine, kidney, gallbladder, adrenal gland, testis, and seminal vesicle among 61 organs in the HPA, at both the protein and RNA levels (Figure S1), consistenting with previous reports [13, 14]. Single-cell sequencing also showed that ACE2 was expressed in only amounts in airway tissue and type II alveolar cells (AT2). In addition, mRNA data of all 64 cell lines in the HPA show low expression of ACE2 in human small-cell lung cancer cells (SCLC-21H, Figure S2). However, ACE2 was not detected at the protein level in airway or lung tissues, indicating that the correlation between protein and transcription levels were widely varied [21]. Thus, limitations arise when predicting protein expression based only on mRNA expression levels.

In a clinical study of 41 patients, it was speculated that SARS-CoV-2 mainly infected the lower respiratory tract [7]. A Chinese research group also isolated this virus from airway EpCs [4, 8]. These results indicate that airway EpCs are possibly a susceptible and primary target for SARS-CoV-2. Furthermore, initial viral entrance may cause cytopathological changes to airway EpCs, resulting in coughing in 76% of patients [7] and further indicating that the lung is also susceptible to virus infection. Thus, the susceptibility of different organs to SARS-CoV-2 is not consistent with corresponding protein expression levels of ACE2 on these organs. We therefore asked (i) why SARS-CoV-2 infects the respiratory tract rather than other organs as its first target organ, (ii) what is the difference between airway EpCs and epithelial-derived cells of other organs, and (iii) whether specific local microenvironment components were required for SARS-CoV-2 infection besides ACE2 that are sufficient to compensate for the low expression of ACE2 in airway EpCs.

To explore these issues, we first outlined the expression profile of virus receptors or receptor-related membrane proteins (collectively named virus microenvironment components) of EpCs located in different organs. We used the human protein atlas (HPA) database [16] to extract the protein expression level of 65 receptors involved in “virus receptor activity” (GO:0001618) of EpCs and epithelial or epithelial-derived cells from 14 organs (16 cell types) (Figure 1A and Supplementary Materials and Methods). By comparing the data of different organs, we found that the expression pattern of these virus microenvironment components varies among the epithelial-derived cells from different organs. We focused on airway EpCs including virus microenvironment components from the nasopharynx and bronchi because these two sites are most likely to be preferred by SARS-CoV-2.

We found serpin family B member 3 (SERPINB3) and cadherin-related family member 3 (CDHR3) were specifically expressed in airway EpCs, which attracted our attention (Figure 1B) for their potential as the viral important microenvironment in the airway. In particular, CDHR3 shows little or no expression within the epithelial-derived cells of other organs (Figure 1C). SERPINB3 is in the serine protease family, and it was reported that the S protein of coronavirus can be activated by transmembrane serine protease family member (TTSP) to complete its membrane fusion process [22, 23]. Thus, the role of SERPINB3 on airway EpCs during SARS-CoV-2 infection merits further study.

Other potential receptors such as xenotropic and polytropic retrovirus receptor 1 (XPR1), in addition to being highly expressed on airway EpCs, are also expressed on the surface of other epithelial-derived cells. XPR1 was also highly expressed on colon gland cells and tonsil squamous EpCs; moderately expressed on EpCs of the oral mucosa, lung, kidney and muscle; and poorly expressed on liver and bile duct cells (Figure 1B, Figure S3). Clinical and epidemiological studies also revealed that diarrhea has been observed in a subset of patients infected with SARS-CoV-2 in addition to the pulmonary manifestations [6–8], indicating that SARS-CoV-2 can also infect the intestinal tract as well as the lung tissue through specific receptors and microenvironment components. This putative intestinal infection require special attention, because it suggests that SARS-CoV-2 also may be transmitted through the fecal-oral route. Stool samples from patients infected with SARS-CoV-2 in the United States also have been positive for the presence of SARS-CoV-2, strongly suggesting fecal transmission [24]. Nucleic acid of the similar coronavirus SARS-CoV was also detected in patients’ stool samples [25–27].

Among the proteins with low expression on airway EpCs, we focused on dystroglycan 1 (DAG1) and found it is only specifically expressed in airway and lung pneumocytes EpCs (Figure 1B). Together these data suggest that the expression levels of putative virus microenvironment components in different types of epithelial-derived cells are organ-specific.

### EpCs from different organs are specific to virus response

We next investigated whether the proteins that respond to SARS-CoV-2 infection display organ-specific expression. We used the same scheme to extract virus-host interaction-related entries (Table S2). In the search category of viral-reactive proteins that modulate host cell morphology or physiology, few were expressed in the liver. Notably, Bcl2-associated agonist of cell death (BAD) and zinc finger C3H-type containing 12A (ZC3H12A) were highly expressed exclusively in airway EpCs; neurotrophic receptor tyrosine kinase 3 (NTRK3) was moderately expressed exclusively in airway EpCs. Another protein, eukaryotic translation initiation factor 2 alpha kinase 4 (EIF1AK4), which was highly expressed in airway EpCs, also was found moderately expressed in the lung, kidney, and muscle (Figure S4).

In terms of viral defense, we found that most of the proteins reactive to viruses are highly expressed in different organs (Figure S4). We found more specific highly expressed proteins in airway EpCs such as calcitonin gene-related peptide-receptor component protein (CRCP), tumor necrosis factor receptor-associated factor 3 interacting protein 1 (TRAF3IP1), adaptor-related protein complex 1 subunit beta 1 (AP1B1), Bcl2, ATP/GTP Binding Protein Like (AGBL)4, AGBL5, adenosine deaminase RNA specific (ADAR), SIN3A, high-temperature requirement A serine peptidase 1 (HTRA1), and ZC3H12A. In addition, several highly expressed proteins were found not only expressed in airway EpCs, but also in the epithelial-derived cells of other organs. For example, apolipoprotein B mRNA editing enzyme catalytic subunit 3A (APOBEC3A) was highly expressed in the colon (glandular cells); transmembrane protein 173 (TMEM173), AP2A2, and ring finger protein 26 (RNF26) were highly expressed in the esophagus (squamous EpCs); RNF26 in the oral mucosa (squamous EpCs); and ribonuclease L (RNASEL) in the tonsil (squamous EpCs). These results indicate that although the responses of different organs to the virus were identical, specific response proteins to the virus vary between different types of EpCs. In particular, highly expressed response proteins on airway EpCs may play a positive role in regulating the propagation and transmission of SARS-CoV-2.

### Interaction network between virus microenvironment components and virus response proteins in airway EpCs

To study whether the virus induces a biological response after infection through airway-specific receptors, we established a network of interactions between the receptors in the airway EpCs and host reactive proteins against the virus. The network controlled several biological processes including virus receptor activity, defense response to virus, viral modulation of host morphology or physiology, detection of virus, evasion or tolerance of host defenses by the virus, viral suppression of host apoptotic process, and viral regulation of viral protein levels in the host cell (Figure 2A). Each protein in the interaction network may, together with the receptor of airway EpCs, be involved in the biological process of host cells after virus infection. For example, Asp-Glu-Ala-Asp (DEAD) box polypeptide 58 (DDX58, also known as retinoic acid-inducible gene I protein, RIG-I), which had the largest number of interacting proteins, acted as a cytoplasmic sensor of viral nucleic acids and played a major role in sensing viral infection and in the activation of a cascade of antiviral responses including the induction of type I interferon and pro-inflammatory cytokines. Interferon induced with helicase C domain 1 (IFIH1), an important paralog gene of DDX58, was involved in the alteration of RNA secondary structure such as translation initiation, nuclear and mitochondrial splicing, and ribosome and spliceosome assembly. DDX58 and IFIH1 were both putative RNA helicases.

**Figure 2.**
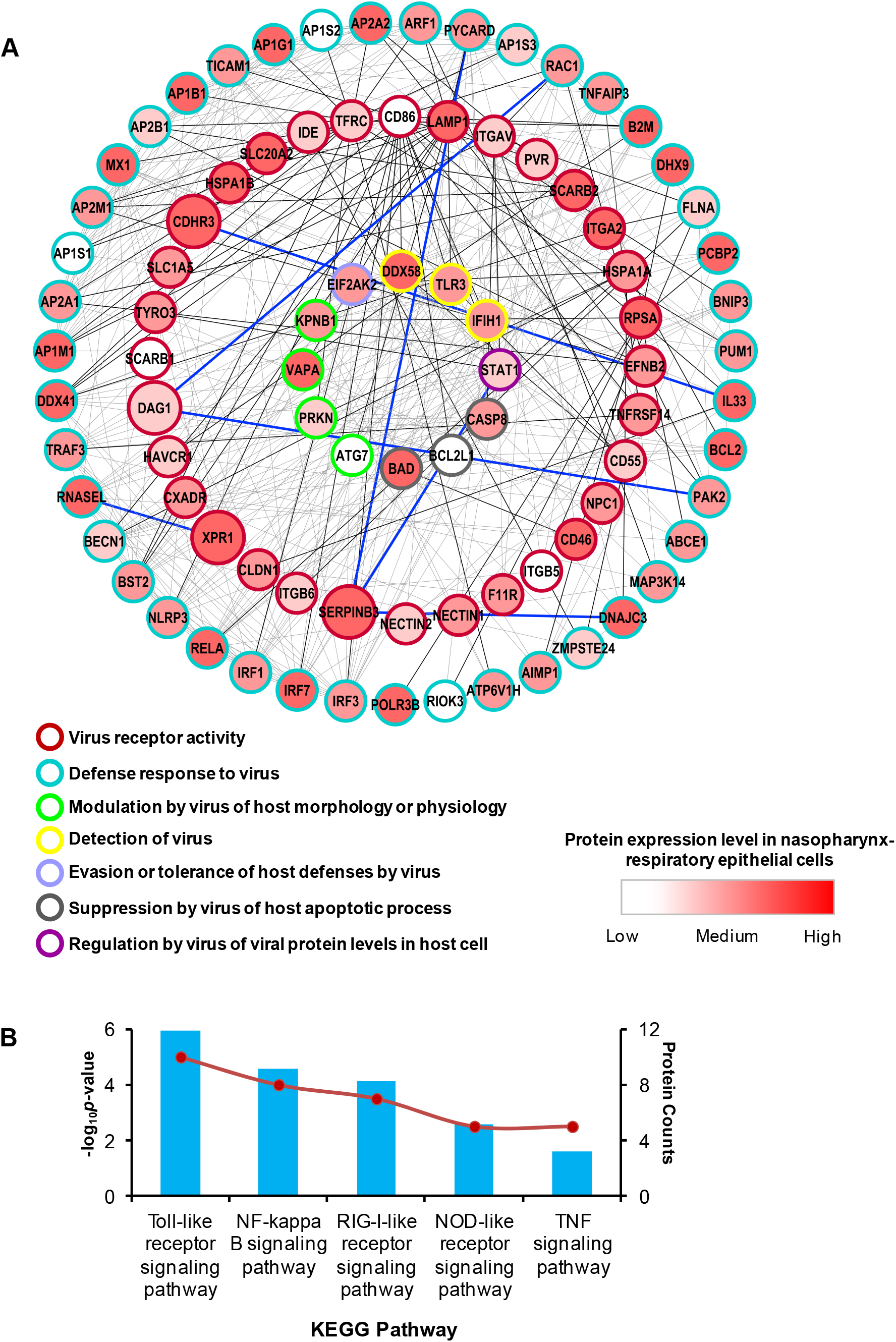
Interaction network between highly expressed virus microenvironment components and virus response proteins in airway epithelial cells (EpCs). (A) Overview of the analysis of the direct interactions between receptors and virus response proteins in airway EpCs. Empty circles with colors represent different functional items. Black and blue lines represent the direct interactions between virus microenvironment components and virus-response proteins in airway EpCs. (B) KEGG pathway analysis of virus microenvironment components and virus response proteins in airway EpCs. Histogram represents log_10_ (*p*-value) of each item on the left. Line chart represents the protein number of each item on the right.

In the interaction network, we focused on the protein interaction of virus microenvironment components with high expression in airway EpCs, which probably perform a specific function in airway EpCs after virus infection. RNASEL, the interacting protein of XPR1, matched these parameters and was specifically highly expressed in airway and tonsil squamous EpCs (Figure 2A). RNASEL probably mediates its antiviral effects through a combination of direct cleavage of single-stranded viral RNAs, inhibition of protein synthesis through the degradation of rRNA, induction of apoptosis, and induction of other antiviral genes. Another receptor protein, DAG1, which was seldom expressed in airway EpCs, interacted with interleukin (IL)-33 (Figure 2A). IL-33 binds to and signals through the IL-1 receptor-like 1 (IL1RL1)/ST2 receptor, which in turn activates nuclear factor (NF)-κB and mitogen-activated protein kinase (MAPK) signaling pathways in target cells.

In addition, we found most of the interacting proteins of virus microenvironment components on airway EpCs are related to the immune response, including toll-like receptors (TLR), NF-κB, RIG-I-like receptor, nucleotide-binding oligomerization domain-containing protein (NOD)-like receptor and TNF signaling pathways (Figure 2B).

Upon detection of a pathogen, epithelia respond by upregulating expression of defensins, which confer immediate protection to the host, and alarmins, which convey danger signals to neighboring EpCs. Epithelia are equipped with TLR surface receptors, which recognize conserved molecular patterns associated with pathogens or tissue damage. TLR signals activate NF-κB through the adaptor protein MyD88 [28]. We found that the most enriched signaling pathways in the interaction network between airway EpC receptors and host-reactive protein are the TLR signaling pathway followed by the NF-κB signaling pathway (Figure 2B). In addition, we found putative activation of the RIG-I-like receptor signaling pathway, which begins a downstream signal pathway to secrete type I interferon in the interaction network. Many proteins related to the production and regulation of type I interferon were also observed in the interaction network (Figure S5). The most enriched item was IL-1β, which is consistent with the fact that the serum of patients infected with SARS-CoV-2 has a higher pro-inflammatory factor IL-1β than that of uninfected people [7]. In addition, the NOD-like receptor and TNF signaling pathways mediate the expression of a variety of inflammatory factors including monocyte chemoattractant protein-1 (MCP-1), interferon-induced protein 10 (IP-10), and TNF-α which are consistent with the COVID-19 clinical characteristics [6, 7].

These results indicate that the virus microenvironment components expressed in airway EpCs, especially those interacting proteins with specific and highly expressed microenvironment components, may be involved in inhibition of host cell biological processes and important immune response pathways related to the antiviral response.

## Conclusion

In this manuscript, we proposes the possibility that airway-specific virus microenvironment components may assist the interaction between ACE2 and SARS-CoV-2, although the expression of ACE2 is very low in the tracheal epithelium that provides other potential directions to study SARS-CoV-2 entry into airway EpCs. Our study also suggests that the clinical symptoms of different organs may be related to the specific expression of virus microenvironment components in these various organs. For example, SARS-CoV-2 might be involved in immune pathways through the interaction network between virus microenvironment components and virus-reactive proteins in airway EpCs, which may be related to the cytokine storm observed in COVID-19 pneumonia patients. These results provide a preliminary research direction for understanding the pathogenesis of SARS-CoV-2 infection in airway EpCs. In addition, our study identified that some virus microenvironment components are not only highly expressed in airway epithelium but are also specifically expressed in other organs, suggesting that more organs may be susceptible to SARS-CoV-2 such as the intestine and kidney.

## Author contribution

LL, JM, WZ, RC and ZW conceived the overall study and designed experiments. LL, JM and LZ performed the data analysis and prepared the figures. LL, JM and RC wrote and edited the manuscript. RC, WL and LZ edited the figures. WZ, ZW and YZ conceived and supervised the data analysis. All authors made important comments to the manuscript.

## References

[1] Q. Li, X. Guan, P. Wu, et al., Early Transmission Dynamics in Wuhan, China, of Novel Coronavirus-Infected Pneumonia, The New England journal of medicine, 382 (2020) 1199–1207.

[2] J.F. Chan, S. Yuan, K.H. Kok, et al., A familial cluster of pneumonia associated with the 2019 novel coronavirus indicating person-to-person transmission: a study of a family cluster, Lancet (London, England), 395 (2020) 514–523.

[3] X. Xu, P. Chen, J. Wang, et al., Evolution of the novel coronavirus from the ongoing Wuhan outbreak and modeling of its spike protein for risk of human transmission, Science China. Life sciences, 63 (2020) 457–460.

[4] P. Zhou, X.L. Yang, X.G. Wang, et al., A pneumonia outbreak associated with a new coronavirus of probable bat origin, Nature, 579 (2020) 270–273.

[5] Y. Ding, H. Wang, H. Shen, et al., The clinical pathology of severe acute respiratory syndrome (SARS): a report from China, The Journal of pathology, 200 (2003) 282–289.

[6] N. Chen, M. Zhou, X. Dong, et al., Epidemiological and clinical characteristics of 99 cases of 2019 novel coronavirus pneumonia in Wuhan, China: a descriptive study, Lancet (London, England), 395 (2020) 507–513.

[7] C. Huang, Y. Wang, X. Li, et al., Clinical features of patients infected with 2019 novel coronavirus in Wuhan, China, Lancet (London, England), 395 (2020) 497–506.

[8] N. Zhu, D. Zhang, W. Wang, et al., A Novel Coronavirus from Patients with Pneumonia in China, 2019, The New England journal of medicine, 382 (2020) 727–733.

[9] J.K. Millet, G.R. Whittaker, Physiological and molecular triggers for SARS-CoV membrane fusion and entry into host cells, Virology, 517 (2018) 3–8.

[10] S.B. Larsen, C.J. Cowley, E. Fuchs, Epithelial cells: liaisons of immunity, Current opinion in immunology, 62 (2019) 45–53.

[11] W. Li, M.J. Moore, N. Vasilieva, et al., Angiotensin-converting enzyme 2 is a functional receptor for the SARS coronavirus, Nature, 426 (2003) 450–454.

[12] X. Zou, K. Chen, J. Zou, et al., Single-cell RNA-seq data analysis on the receptor ACE2 expression reveals the potential risk of different human organs vulnerable to 2019-nCoV infection, Frontiers of medicine, (2020) doi: 10.1007/s11684-11020-10754-11680.

[13] F. Qi, S. Qian, S. Zhang, et al., Single cell RNA sequencing of 13 human tissues identify cell types and receptors of human coronaviruses, Biochemical and biophysical research communications, (2020) pii: S0006-0291X(0020)30523-30524.

[14] W. Lin, L. Hu, Y. Zhang, et al., Single-cell Analysis of ACE2 Expression in Human Kidneys and Bladders Reveals a Potential Route of 2019-nCoV Infection, bioRxiv, (2020) 2020.2002.2008.939892.

[15] F. Hikmet, L. Méar, M. Uhlén, et al., The protein expression profile of ACE2 in human tissues, bioRxiv, (2020) 2020.2003.2031.016048.

[16] M. Uhlen, L. Fagerberg, B.M. Hallstrom, et al., Tissue-based map of the human proteome, Science (New York, N.Y.), 347 (2015) 1260419.

[17] T.G.O. Consortium, The Gene Ontology Resource: 20 years and still GOing strong, Nucleic acids research, 47 (2019) D330–d338.

[18] T.U. Consortium, UniProt: the universal protein knowledgebase, Nucleic acids research, 45 (2017) D158–d169.

[19] D. Szklarczyk, J.H. Morris, H. Cook, et al., The STRING database in 2017: quality-controlled protein-protein association networks, made broadly accessible, Nucleic acids research, 45 (2017) D362–D368.

[20] P. Shannon, A. Markiel, O. Ozier, et al., Cytoscape: a software environment for integrated models of biomolecular interaction networks, Genome research, 13 (2003) 2498–2504.

[21] D.P. Nusinow, J. Szpyt, M. Ghandi, et al., Quantitative Proteomics of the Cancer Cell Line Encyclopedia, Cell, 180 (2020) 387–402.e316.

[22] S. Bertram, R. Dijkman, M. Habjan, et al., TMPRSS2 activates the human coronavirus 229E for cathepsin-independent host cell entry and is expressed in viral target cells in the respiratory epithelium, Journal of virology, 87 (2013) 6150–6160.

[23] S. Gierer, S. Bertram, F. Kaup, et al., The spike protein of the emerging betacoronavirus EMC uses a novel coronavirus receptor for entry, can be activated by TMPRSS2, and is targeted by neutralizing antibodies, Journal of virology, 87 (2013) 5502–5511.

[24] M.L. Holshue, C. DeBolt, S. Lindquist, et al., First Case of 2019 Novel Coronavirus in the United States, The New England journal of medicine, 382 (2020) 929–936.

[25] W.K. Leung, K.F. To, P.K. Chan, et al., Enteric involvement of severe acute respiratory syndrome-associated coronavirus infection, Gastroenterology, 125 (2003) 1011–1017.

[26] J.S. Peiris, C.M. Chu, V.C. Cheng, et al., Clinical progression and viral load in a community outbreak of coronavirus-associated SARS pneumonia: a prospective study, Lancet (London, England), 361 (2003) 1767–1772.

[27] J.S. Peiris, S.T. Lai, L.L. Poon, et al., Coronavirus as a possible cause of severe acute respiratory syndrome, Lancet (London, England), 361 (2003) 1319–1325.

[28] A.E. Price, K. Shamardani, K.A. Lugo, et al., A Map of Toll-like Receptor Expression in the Intestinal Epithelium Reveals Distinct Spatial, Cell Type-Specific, and Temporal Patterns, Immunity, 49 (2018) 560–575.e566.

