## Supplementary Figures and Tables for "Potential microenvironment of SARS-CoV-2 infection in airway epithelial cells revealed by Human Protein Atlas database analysis"

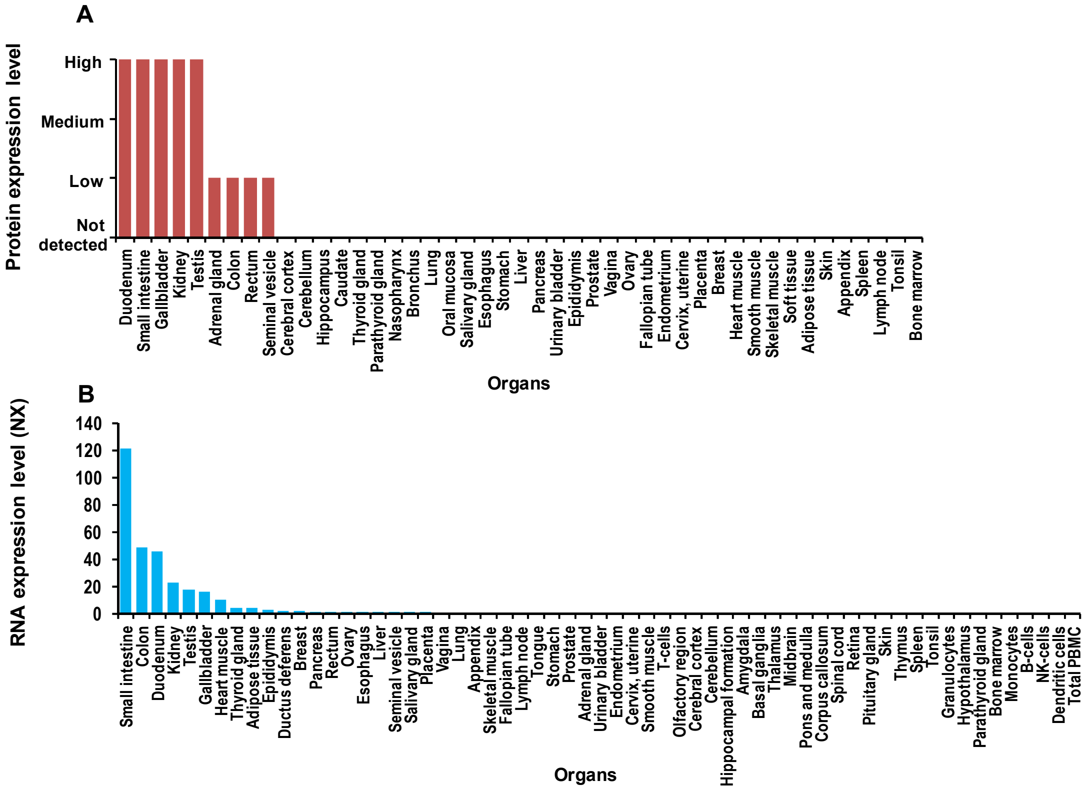


**Figure S1. Expression of angiotensin converting enzyme 2 (ACE2) in different organs.** The expression datasets of protein levels (A) and mRNA levels (B) of ACE2 were all downloaded from HPA. (https://www.proteinatlas.org/ENSG00000130234-ACE2/tissue).

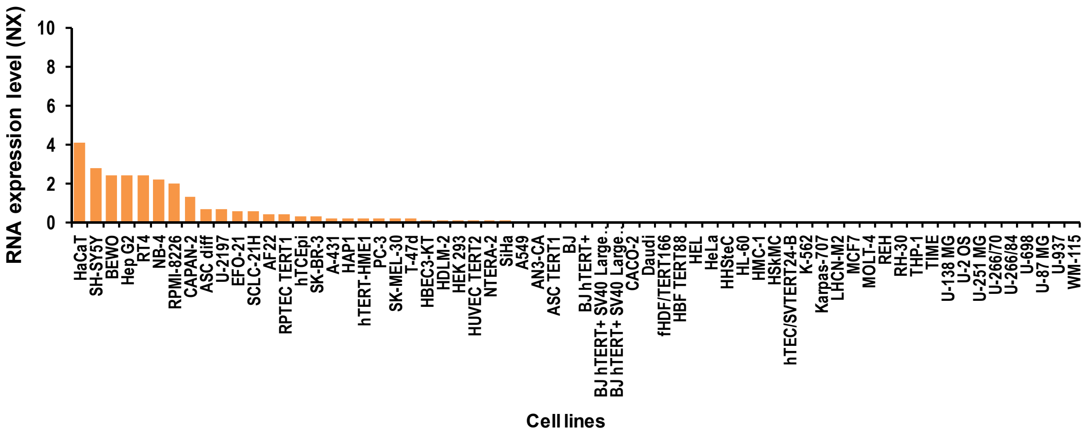


**Figure S2. Expression of angiotensin converting enzyme 2 (ACE2) in different cell lines.** The mRNA expression dataset was downloaded from HPA (https://www.proteinatlas.org/ENSG00000130234-ACE2/cell).

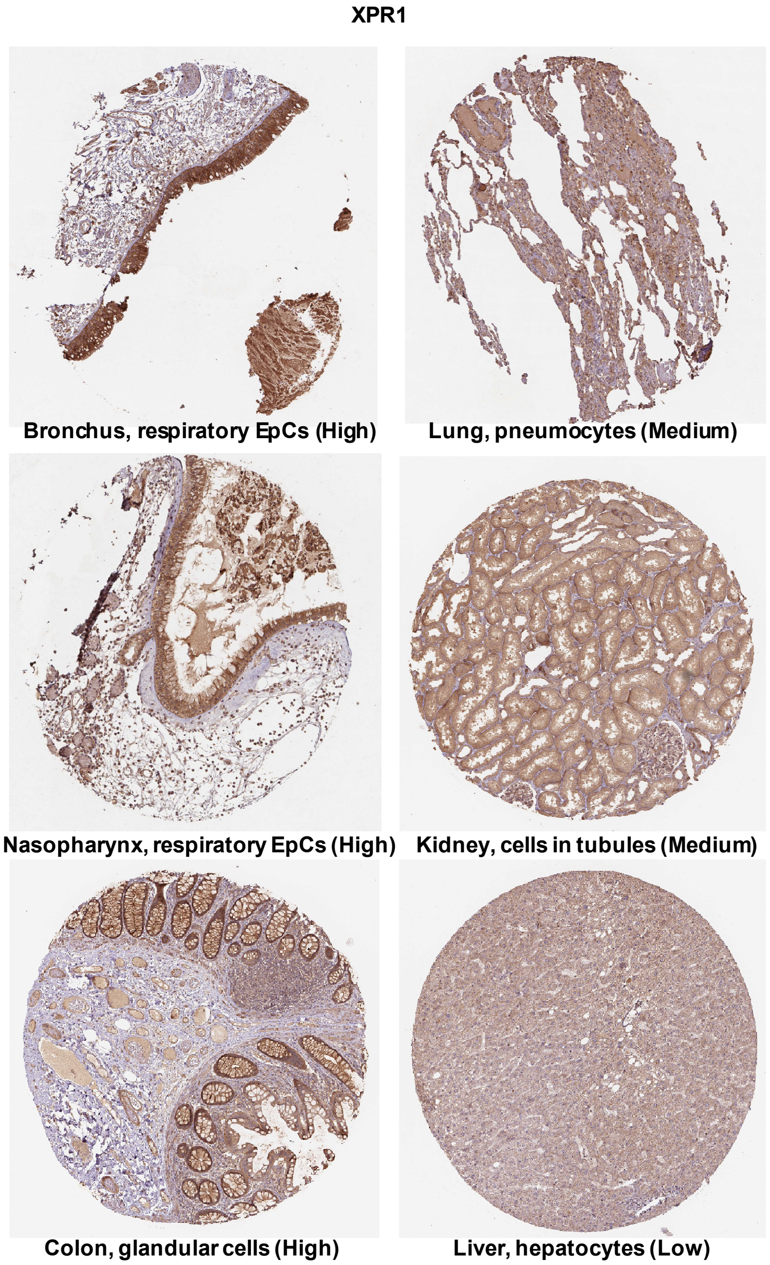


**Figure S3. Immunohistochemistry analysis of XPR1 in epithelial or epithelial-derived cells of different organs.** The staining pictures are downloaded from "The Tissue Atlas" of HPA (https://www.proteinatlas.org/ENSG00000143324-XPR1/tissue).

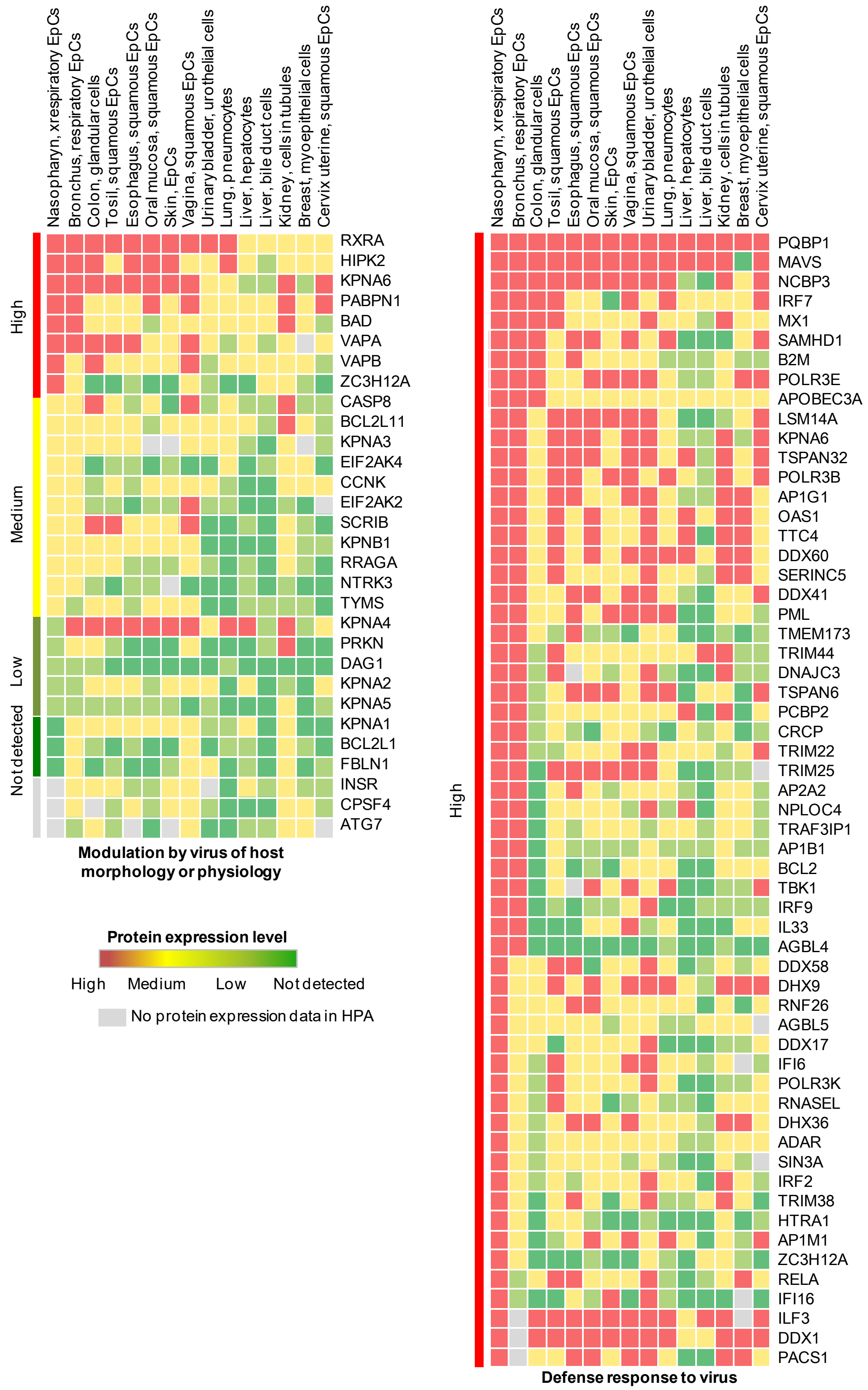


**Figure S4. Heat map analysis of virus-response proteins in epithelial-derived cells from different organs.** The boxes with red, yellow, green, and dark green represent the expression levels of receptors with high, medium, low, and not detected values, respectively.

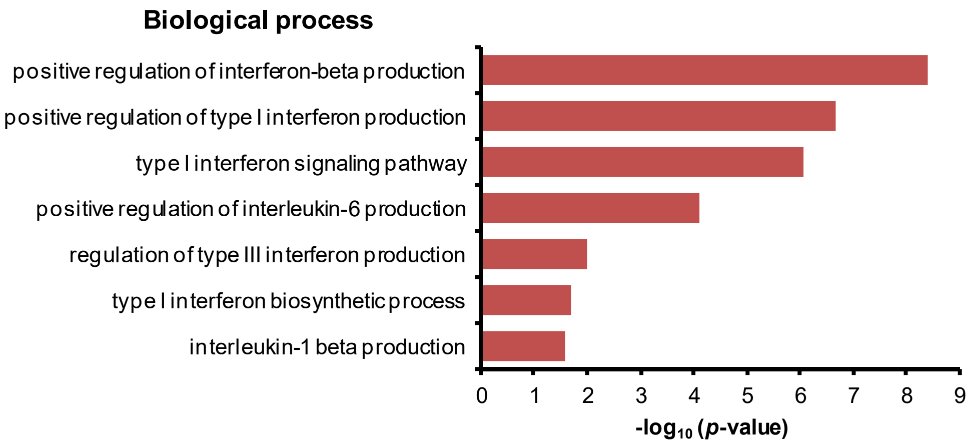


**Figure S5. Biological processes of interleukin (IL)-1β production of virus microenvironment components and virus response proteins in airway EpCs.**

### Supplementary tables

#### Table S1. Expression profile of virus microenvironment components in "virus receptor activity" in different human tissues in the Human Protein Atlas.

| Tissue/ Cell Type | Nasopharynx | Bronchus | Lung | Colon | Tonsil | Esophagus | Oral mucosa | Skin | | Urinary bladder | Vagina | Liver | | kidney | Breast | Cervix, uterine |
|---|---|---|---|---|---|---|---|---|---|---|---|---|---|---|---|---|
| Gene Name | Respiratory EpCs | Respiratory EpCs | Pneumo -cytes | Glandular cells | Squamous EpCs | Squamous EpCs | Squamous EpCS | Keratino -cytes | EpCs | Urothelial cells | Squamous EpCS | Hepato -cytes | Bile duct cells | Cells in tubules | Myoepithelial cells | Squamous EpCs |
| ACE2 | None | None | None | Low | None | None | None | None | None | None | None | None | None | High | None | None |
| TFRC | Low | Low | None | Medium | Medium | Medium | Medium | Low | Medium | Medium | Medium | Low | None | Medium | Low | Low |
| LDLR | None | None | None | Medium | None | None | None | None | None | Medium | None | Medium | None | Medium | None | None |
| EFNB3 | None | None | Low | None | None | None | None | None | None | N/A | None | None | None | None | None | N/A |
| CD86 | None | Low | None | Medium | N/A | Low | Low | None | Medium | Medium | Low | None | N/A | None | None | None |
| XPR1 | High | High | Medium | High | High | Medium | Low | Low | Medium | Medium | Medium | Low | Low | Medium | Medium | Medium |
| HTR2A | None | None | None | None | None | None | None | None | None | None | None | None | None | None | None | None |
| SERPINB3 | High | High | None | None | None | Medium | Low | Low | Low | Medium | Low | None | None | None | None | Low |
| ITGAV | Low | Medium | None | Medium | High | Medium | Medium | Medium | Medium | High | None | None | Medium | High | Low | Low |
| SLC10A1 | None | None | None | None | None | None | None | None | None | None | None | Medium | None | None | None | None |
| CD81 | None | None | None | None | None | None | None | Low | None | None | None | None | None | None | None | None |
| CR1 | None | None | None | None | None | None | None | None | None | None | None | None | None | None | None | None |
| NCAM1 | None | None | None | None | None | None | None | None | None | None | None | None | None | None | None | N/A |
| GPR15 | None | None | None | None | None | None | None | None | None | None | None | Low | None | None | None | None |
| NPC1 | Medium | Medium | Medium | Medium | Medium | Medium | Medium | Low | Medium | Medium | Medium | Medium | Low | High | Low | Low |
| HSPA1A | Medium | High | Medium | Medium | Medium | High | Medium | Medium | Medium | High | High | Medium | Medium | Medium | Medium | High |
| SLC20A2 | High | High | None | High | Low | None | None | Low | None | High | None | None | High | Medium | None | None |
| CLEC4G | None | None | None | None | None | None | None | None | None | None | None | None | None | None | None | None |
| CDHR3 | High | High | None | None | None | None | None | None | None | None | None | None | None | None | None | None |

| | | | | | | | | | | | | | | | | |
|---|---|---|---|---|---|---|---|---|---|---|---|---|---|---|---|---|
| AXL | None | None | Medium | Medium | None | None | None | None | None | Low | None | Low | None | Medium | None | None |
| LAMP1 | High | High | High | High | High | High | High | Medium | High | High | High | High | High | High | Medium | High |
| CDK1 | None | None | None | Medium | Medium | Medium | Medium | None | Medium | Medium | Medium | None | None | Medium | None | Low |
| SLC1A5 | Medium | Medium | Low | Medium | Medium | Medium | Medium | Low | Medium | High | Medium | Low | None | Medium | Medium | Low |
| ANPEP | N/A | None | None | Medium | None | None | None | None | None | None | None | High | High | High | None | Low |
| CD4 | None | None | None | None | None | None | None | None | None | None | None | None | None | None | None | None |
| CLEC4M | None | None | None | None | None | None | None | None | None | None | None | None | None | None | None | None |
| IDE | Low | Medium | None | Medium | Low | Low | Low | Low | Medium | Low | None | Low | Low | Medium | Medium | Low |
| CXADR | Medium | Medium | None | Medium | Medium | Low | Low | Medium | Medium | High | Medium | Medium | Medium | Low | Low | Medium |
| SELPLG | N/A | None | None | None | None | None | None | None | None | None | None | None | None | None | None | None |
| CLDN1 | Medium | Medium | None | None | High | Low | Low | Medium | Medium | Medium | Medium | Medium | Medium | Medium | Low | Medium |
| CD80 | None | None | None | None | None | None | None | None | None | None | None | None | None | None | None | None |
| NECTIN4 | None | None | None | None | Medium | Medium | Medium | Medium | Medium | Medium | Low | None | None | Low | Medium | Low |
| TNFRSF4 | None | None | None | None | None | None | None | None | None | None | None | None | None | None | None | None |
| CR2 | None | None | None | None | None | None | None | None | None | None | None | None | None | None | None | None |
| EFNB2 | Medium | Medium | Medium | Medium | Medium | Medium | Low | Medium | Medium | Low | Medium | Medium | Low | Medium | Medium | Medium |
| MOG | None | None | None | None | None | None | None | None | None | None | None | None | None | None | None | None |
| NECTIN2 | Low | Low | Medium | Medium | Medium | Low | None | Low | Low | Low | Low | Medium | Medium | Medium | None | None |
| NECTIN1 | Medium | Medium | High | High | Medium | Medium | Medium | Low | High | Medium | Medium | Low | High | Medium | Medium | Medium |
| GYPA | None | None | None | None | None | None | None | None | None | None | None | None | None | None | None | None |
| DAG1 | Low | Low | Low | Low | None | None | None | Low | None | None | None | None | None | None | None | None |
| SLAMF1 | N/A | None | None | None | None | None | None | None | None | None | None | None | None | None | None | None |
| ICAM1 | None | None | High | None | Medium | None | None | None | None | Low | None | None | None | None | None | None |
| EGFR | N/A | None | None | None | Low | None | None | None | None | Low | Low | Low | None | None | None | N/A |
| ITGB5 | None | Low | Low | Low | Medium | None | Low | Medium | None | Low | None | Medium | Medium | Medium | Medium | None |

| | | | | | | | | | | | | | | | |
|---|---|---|---|---|---|---|---|---|---|---|---|---|---|---|---|
| ITGB1 | None | None | None | Low | None | None | None | None | None | Low | None | None | None | None | Low | None |
| ITGB3 | None | None | None | None | None | None | None | None | None | None | None | None | None | None | None | None |
| HSPA1B | High | High | None | Low | Medium | High | Medium | High | High | High | Medium | Low | Medium | None | None | High |
| TNFRSF14 | Medium | Medium | None | Medium | Low | Low | None | Low | Low | High | Low | Low | Medium | High | None | Low |
| SLC52A2 | Low | Low | None | Medium | None | Low | Low | Low | None | Medium | None | None | None | Low | None | Low |
| SCARB1 | None | Medium | None | None | None | None | None | None | None | None | None | None | None | None | None | None |
| CD55 | Low | Low | High | Low | Low | None | None | Low | Low | Low | Low | None | None | Low | None | None |
| ITGA5 | None | None | None | None | None | None | None | None | None | None | None | None | None | None | None | None |
| RPSA | High | High | Medium | High | High | High | High | High | High | High | High | Medium | None | Medium | High | N/A |
| SCARB2 | High | High | Medium | High | Medium | Low | Low | High | Medium | High | None | High | Medium | Medium | None | None |
| ITGA2 | High | High | Medium | High | High | Medium | Medium | Medium | Medium | High | Medium | Low | Medium | High | High | Medium |
| CD46 | High | Medium | Medium | High | Low | Low | Low | None | Low | High | Low | Medium | Low | Medium | High | Low |
| HAVCR1 | Low | Low | None | Medium | Low | None | None | Low | Low | Medium | Low | None | Medium | Medium | None | Low |
| HYAL2 | None | None | None | None | None | None | None | None | None | Low | None | None | None | High | Low | N/A |
| CD209 | None | None | None | None | None | None | None | None | None | None | None | None | None | None | None | N/A |
| PVR | Low | Low | None | Medium | Low | Low | None | None | None | None | None | None | Low | Low | Low | None |
| ITGB6 | Low | None | Medium | Medium | Medium | None | None | None | None | High | None | None | None | Low | None | None |
| MRC1 | None | None | None | Low | None | None | N/A | None | None | None | None | None | None | Low | None | None |
| F11R | Medium | Medium | Low | Medium | Medium | Medium | Medium | Medium | Medium | High | Medium | Medium | Medium | Medium | None | Medium |
| TYRO3 | Medium | Medium | None | Medium | Medium | Medium | Low | Low | Medium | Medium | Low | Low | Low | Medium | Low | Medium |
| DPP4 | None | None | None | Low | None | None | None | None | None | None | None | Low | None | High | None | None |

None: Not detected.

N/A: No protein expression data in HPA.

**Table S2. Detailed lists of GO terms used in this study.**

| ID | Name Space | Name |
|---|---|---|
| GO:0001618 | Molecular function | virus receptor activity |
| GO:0002230 | Biological process | positive regulation of defense response to virus by host |
| GO:0006948 | Biological process | induction by virus of host cell-cell fusion |
| GO:0009597 | Biological process | detection of virus |
| GO:0009615 | Biological process | response to virus |
| GO:0019045 | Biological process | latent virus replication |
| GO:0019048 | Biological process | modulation by virus of host morphology or physiology |
| GO:0019049 | Biological process | evasion or tolerance of host defenses by virus |
| GO:0019050 | Biological process | suppression by virus of host apoptotic process |
| GO:0019061 | Biological process | uncoating of virus |
| GO:0019064 | Biological process | fusion of virus membrane with host plasma membrane |
| GO:0019075 | Biological process | virus maturation |
| GO:0030683 | Biological process | evasion or tolerance by virus of host immune response |
| GO:0039502 | Biological process | suppression by virus of host type I interferon-mediated signaling pathway |
| GO:0039526 | Biological process | modulation by virus of host apoptotic process |
| GO:0039562 | Biological process | suppression by virus of host STAT activity |
| GO:0039656 | Biological process | modulation by virus of host gene expression |
| GO:0046719 | Biological process | regulation by virus of viral protein levels in host cell |
| GO:0046725 | Biological process | negative regulation by virus of viral protein levels in host cell |
| GO:0046726 | Biological process | positive regulation by virus of viral protein levels in host cell |
| GO:0046778 | Biological process | modification by virus of host mRNA processing |
| GO:0050687 | Biological process | negative regulation of defense response to virus |
| GO:0050688 | Biological process | regulation of defense response to virus |
| GO:0050689 | Biological process | negative regulation of defense response to virus by host |
| GO:0050690 | Biological process | regulation of defense response to virus by virus |
| GO:0051607 | Biological process | defense response to virus |
